# Thermopriming prewires orchestrated regulatory programs to enhance heat stress adaptability

**DOI:** 10.64898/2026.11.12.766148

**Authors:** Akshay U Nair, Shubham Vishwakarma, Rajashekar Varma Kadumuri, Felix B. Fritschi, Sreenivas Chavali, Annapurna Devi Allu

## Abstract

- Plants can remember earlier stress exposures (priming), which improves tolerance to later, more severe stress. How this stress memory is organised in crops remains largely unresolved.
- We studied five-day-old rice seedlings subjected to severe heat stress with or without thermopriming, combining phenotyping, time-resolved transcriptomic and epigenomic profiling with systems biology approaches and targeted molecular experiments.
- Compared with unprimed seedlings, primed seedlings showed a restrained, lower-entropy transcriptional response spanning diverse biological processes. This response was driven by fewer transcription factors that modulate regulons for both stress response and growth, together contributing to better adaptability to severe stress.
- Primed seedlings maintained global H3K27me3 levels, which declined in unprimed seedlings, linking retention rather than removal of this mark to the restrained response.
- Priming led to the accumulation of memory chaperone transcripts, enabling a rapid rise in protein abundance under severe stress. This supports better post-translational regulation, alongside the coordinated transcriptional response.
- Overall, thermopriming establishes pre-wired programmes that enhance adaptability through regulatory order and speed, not amplitude, and repressive-mark retention, not removal, during early development. These findings have implications for (i) understanding heat stress memory in addressing climate change driven heat extremes, and (ii) engineering climate-resilient crops.

## Introduction

Global warming poses a serious threat to crop productivity, as high temperatures adversely affect key developmental stages, disrupt physiological processes and ultimately result in substantial yield loss (Xu *et al*., 2020; Mthiyane *et al*., 2024; Singh *et al*., 2024). To adapt to rapidly fluctuating and challenging environments, plants have the ability to ‘remember’ prior stress encounters and use that ‘memory’ to better acclimate to future stresses. This adaptive strategy, known as ‘priming,’ provides acquired tolerance for subsequent stresses (Lämke & Bäurle, 2017; Liu *et al*., 2022; Nair *et al*., 2022). The magnitude, period and frequency of priming stimuli can influence the duration and nature of stress memory, which can range from short-to long-term somatic memory or transgenerational memory (Sani *et al*., 2013; Wang *et al*., 2016; Bäurle & Trindade, 2020).

Acquired thermotolerance, in which a sub-lethal heat stress serves as a priming cue, enhances tolerance to subsequent heat stress (Charng *et al*., 2007; Lämke *et al*., 2016; Serrano *et al*., 2019). Individual molecular-level studies, predominantly in the model plant *Arabidopsis thaliana*, show that (i) thermopriming enhances the transcription of key heat shock factors (*HSFs*) and other stress-responsive genes and (ii) such HSFs collaborate with chromatin remodelers or histone-modifying enzymes to alter chromatin accessibility and establish permissive epigenetic states that contribute to heat stress adaptability (Brzezinka et al., 2016; Lämke *et al*., 2016; Liu *et al*., 2018; Friedrich *et al*., 2021; Yamaguchi *et al*., 2021). Current models of thermomemory, however, rest majorly on single memory genes, whose expression is maintained or re-induced more strongly when stress recurs, with the help of HSFs and chromatin modifiers such as REF6, an H3K27me3 demethylase (Brzezinka et al., 2016; Lämke *et al*., 2016; Yamaguchi & Ito, 2021; Yamaguchi *et al*., 2021). How dynamic transcriptional reprogramming after priming shapes the response to repeated stress, in terms of scale, coordination, and regulation, and which transcriptional and epigenetic features drive it, is still largely unclear. This is particularly pertinent for rice and other crops, where thermomemory is poorly characterised.

To address this, here we carried out extensive experiments and integrated time-resolved, genome-wide transcriptomic and epigenetic profiles in rice, a staple crop. We show that thermopriming pre-wires a smaller, more ordered transcriptional response to a subsequent triggering heat stress, characterised by fewer transcription factors drawn from smaller paralog families, coordinated regulatory networks, retention of H3K27me3, and higher levels of memory chaperone.

## Materials and Methods

### Plant materials and growth conditions

Rice seedlings (*Oryza sativa* subsp. indica cv. Nagina22) were cultivated hydroponically using 1X Hoagland nutrient solution, following the formulation of Hoagland & Arnon (1950) with slight modifications. Before germination, seeds were surface sterilised using 1% (v/v) sodium hypochlorite for 10 min and subsequently rinsed several times with sterile distilled water. Around 15-20 sterilised seeds were sown in each phyta-jar (transparent container with an aerated lid and valve). The nutrient solution volume was adjusted between 40 and 80 mL per jar based on seedling growth stage. Plants were grown under controlled conditions in a growth chamber with a 14 h light/10 h dark cycle, maintaining day-time and night-time temperatures of 28 °C and 26 °C, respectively. The relative humidity was set at 65-70%.

### Stress regime and physiological measurements

Heat stress experiments were conducted on 5-day-old rice seedlings according to a previously established protocol (Nair *et al*., 2026). Control plants were maintained under optimal growth conditions at 28 °C. For priming, seedlings were exposed to a sub-lethal heat treatment at 37 °C for 2 h, followed by a 2 h recovery at 28 °C. Thereafter, both primed and unprimed seedlings were subjected to a lethal heat stress (triggering treatment) at 45 °C for 4 h. Unprimed seedlings received only the triggering stimulus without the priming step. Growth and physiological traits, including shoot length, root length, and fresh weight, were measured on the 8^th^ day following heat stress. Leaf surface temperature was recorded immediately after heat exposure using a thermal infrared camera (FLIR A325sc), and data were analysed using FLIR RIRmax software.

### Transcriptome analysis

Shoot tissues from five-day-old rice seedlings were harvested at distinct time points: immediately after priming, before the triggering stress, and immediately following 4 h of triggering stress (45 °C) in primed and unprimed seedlings for three biological replicates. Total RNA was extracted from 100 mg of tissue using the PureLink™ RNA Mini Kit (Invitrogen), and RNA quality and integrity were assessed with the Agilent 2100 Bioanalyzer. Approximately 2 μg of high-quality RNA was used to prepare cDNA libraries using the Illumina HiSeq2000 RNA Library Preparation Kit, followed by paired-end sequencing. Raw reads underwent quality control with FASTQC, and adapter trimming was performed using Trim Galore. Clean reads were aligned to the *Oryza_sativa*_aus_N22.assembly reference genome (from Rice Genome Hub) using HISAT2 (Kim *et al*., 2019), and gene-level read counts were obtained with HTSeq (Anders *et al*., 2015). Differential expression analysis was performed using DESeq2 (Love *et al.,* 2014), and significantly upregulated (Log2FC ≥ 1.5) and downregulated (Log2FC ≤ −1.5) genes were selected based on an adjusted P-value < 0.05.

### Gene expression dispersion entropy

The distribution of Log_2_ fold change values for upregulated and downregulated differentially expressed genes (DEGs) was visualised using box plots, with median values indicated. Statistical differences between the two distributions were assessed using the Wilcoxon rank-sum test. To quantify gene expression dispersion at a single time point, entropy (*H_stp_*) was calculated based on transcriptome-wide variance using the equation:

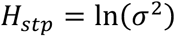

The variance (σ^2^) of the whole-transcriptome DE distribution is computed using the “var” function in R (v3.6.2). Similar calculations were simulated for DEG TFs as well.

### Network entropy analysis

To calculate the approximate von Neumann entropy of a directed graph, the following equation is used

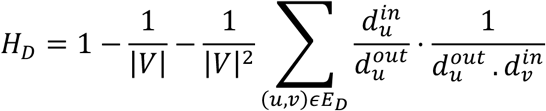

For a graph G, *E_D_* is the directed edge set and node set V. The edge commences at node u and ends at node 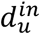 is the in-degree of node u, 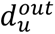 is the out-degree of node u. Shannon entropy was also calculated for the static directed network in After-triggering time point using the equation.

### Identification of transcription factors and TF-target interactions

Transcription factors (TFs) among the differentially expressed genes (DEGs) were identified by submitting their nucleotide sequences to the iTAK database (Zheng *et al*., 2016). To assess family-wise enrichment, the number of DEG TFs in each TF family was normalised by the total number of known TFs in that family in rice. To predict regulatory interactions between TFs and their targets, we used the Regulation Prediction tool from the PlantRegMap database (Tian *et al*., 2020). Promoter sequences spanning 1 kb upstream of DEG transcription start sites were retrieved from RAP-DB (Sakai *et al*., 2013). The database scans these promoter regions for known TF binding motifs to infer putative TF-target interactions and identifies overrepresented TFs likely regulating the DEGs. DEG TFs with enriched binding sites in the 1 kb promoter regions were retained for further analysis. Predicted TF-target pairs were then filtered based on co-expression analysis. Pearson correlation coefficients were computed using FPKM values across biological replicates, separately for control-primed and control-unprimed conditions. TF-target correlations with FDR-adjusted P < 0.05 were considered significant.

### TF-Target network analyses and module detection

TF-target interaction pairs identified under unprimed and primed conditions were used to construct regulatory networks using Cytoscape v3.9.1. Network topology metrics, including indegree and outdegree, were calculated using the Network Analysis tool in Cytoscape. To identify functional clusters, TF–target networks were partitioned into modules using the GLay community clustering algorithm implemented in the clusterMaker2 plugin (Shannon *et al*., 2003; Morris *et al*., 2011). Gene Ontology (GO) enrichment analysis for genes within each module was conducted using the GO-Slim function of the PANTHER database (Thomas *et al*., 2022). The number of TFs and the percentage of known heat stress-responsive genes in each module were determined using STIFDB v.2 (Naika *et al*., 2013), a curated database of stress-responsive transcription factors in plants.

### DEG-Paralog cluster analysis

Differentially expressed transcription factors (DEG TFs) identified under unprimed and primed conditions were queried in the Plant Ensembl database to retrieve their corresponding paralogous genes. The dplyr package in R was used to compute the size of each DEG TF’s paralog cluster. Paralog clusters were categorised into five size-based groups: 1, 2, 3, 4, and ≥5 members. Statistical significance of the distribution differences between conditions was assessed using Fisher’s exact test.

### Gene Ontology and protein class enrichment analysis

Gene Ontology (GO) enrichment analyses were performed for the DEGs and TF-target modules using ShinyGO (v0.76; Ge *et al*., 2020). To identify enriched protein classes, the gene sets were further analysed using the PANTHER classification system (https://pantherdb.org/).

### Total protein extraction

Total protein was extracted from shoot tissue of 5-day-old rice seedlings using the Pierce™ Plant Total Protein Extraction Kit (Cat. No. A44056). Shoot tissue frozen in liquid nitrogen was ground to a fine powder, and 150 mg of the powdered tissue was transferred to the provided cartridge. A volume of 100 μL of denaturing lysis buffer was added, and the tissue was further homogenised using the plastic grinding rod (∼60 strokes). The cartridge was capped and incubated at room temperature for 1–2 min. Lysates were centrifuged at 16,000 x g for 5 min, and the supernatant containing denatured total protein was collected into a microcentrifuge tube and stored at -80 °C for further use.

### Histone protein extraction

Histone proteins were extracted from shoot tissues of 5-day-old rice seedlings using the EpiQuik™ Total Histone Extraction Kit (Cat. No. OP-0006-100). Approximately 200 mg of frozen shoot tissue was homogenised in 1 mL of 1X pre-lysis buffer by vigorous vortexing, with intermittent cooling on ice. The homogenate was transferred to a 2 mL microcentrifuge tube and centrifuged at 10,000 rpm for 1 min at 4 °C. The supernatant was discarded, and the pellet was resuspended in 600 µL of lysis buffer, followed by incubation on ice for 30 min. The lysate was centrifuged at 12,000 rpm for 5 min at 4 °C, and the clear supernatant was collected. A Balance-DTT buffer was prepared by diluting DTT in Balance buffer at a 1:500 ratio, and 180 µL of this buffer was added to the protein extract. The final histone protein samples were stored at - 80 °C for further use.

### Immunoblot analysis

Total protein extracts were used to quantify heat shock proteins (HSPs) at early and later time points, while histone extracts were used to assess histone modifications at corresponding time intervals. Western blotting was performed using the following primary antibodies: anti-HSP101 (ab80121; Abcam), anti-HSP21 (AS08 285; Agrisera), anti-Tubulin beta chain (AS10 680; Agrisera), anti-H3 (ab1791; Abcam), anti-H3K4me3 (ab8580; Abcam), anti-H3K27me3 (ab6002; Abcam), anti-H3K27ac (ab4729; Abcam), and anti-H2A.Z (ab4174; Abcam). Signal detection was carried out using the Pierce™ ECL Western Blotting HRP Substrate (Thermo Fisher Scientific). Band intensities corresponding to specific proteins or histone modifications were quantified using ImageJ software (Schneider *et al*., 2012). Each antibody-based assay was performed with three independent biological replicates.

### Quantitative Real-Time PCR (qRT-PCR)

Total RNA (2 μg) was treated with DNase using the Turbo DNase Kit (Thermo Fisher Scientific) to remove any genomic DNA contamination. RNA concentration and purity were assessed using a NanoDrop™ One/OneC Microvolume UV-Vis Spectrophotometer (Thermo Scientific). First-strand cDNA synthesis was performed using the RevertAid H Minus First Strand cDNA Synthesis Kit (Thermo Scientific). Gene-specific primers for quantitative real-time PCR (qRT-PCR) were designed using QUANTPRIME, a primer design tool optimised for high-throughput qPCR (Arvidsson *et al*., 2008). Expression analysis of candidate genes was conducted using the SYBR Green assay (POWER SYBR® GREEN PCR Master Mix) on BIO-RAD CFX384 and CFX96 Real-Time PCR Systems with gene-specific forward and reverse primers. Melt curve analysis was included to verify primer specificity and amplification efficiency. The housekeeping gene *OsACTIN11* was used as an internal reference for normalising gene expression across samples. Relative expression levels were calculated by converting Ct values into Log₂ fold change (Log2FC) values. A complete list of primers used in this study is provided in the supplementary table (**Table S1.8**).

### Chromatin Immunoprecipitation (ChIP) and library preparation

For ChIP sequencing experiment five-day-old rice seedlings were used. Immediately following 4 h of triggering stress (45 °C) in both primed and unprimed seedlings, about 2 g of shoot tissue was fixed in 1% formaldehyde. Nuclei isolation and chromatin extraction were performed according to the protocols and extraction buffers described by Song *et al*. (2016) and Ramirez-Prado *et al*. (2021). Chromatin was sheared using a Sonics Vibra-Cell sonicator to obtain DNA fragments of 300-500 bp. Sonication efficiency and fragment size distribution were verified using a TapeStation system (Agilent). For each sample, 40 µL of Protein G Dynabeads (Thermo Fisher Scientific) were washed with PBS-BSA and incubated overnight at 4 °C with 4 µg of anti-H3K27me3 antibody (ab6002; Abcam). The sheared nuclear lysate was diluted four-fold and incubated overnight with the antibody-bound beads. The beads were then collected on a magnetic stand and washed sequentially with low-salt, high-salt, LiCl and TE buffers. Immunocomplexes were eluted using the elution buffer described by Ramirez-Prado *et al*. (2021). Cross-links were reversed by incubation overnight at 65 °C, followed by treatment with RNase A and proteinase K. DNA was purified using the QIAquick PCR Purification Kit (Qiagen). The immunoprecipitated DNA was used to prepare ChIP-seq libraries with the Illumina-compatible NEXTflex Rapid DNA-Seq Kit 2.0 (Revvity), following the manufacturer’s instructions.

### ChIP-Sequencing analysis

Paired-end reads were trimmed with Trim Galore (v0.6.10; Phred < 20), aligned to the N22 genome with BWA-MEM (v0.7.18), and duplicates were marked with Picard (v3.3.0). H3K27me3 peaks were called for each replicate against its input using MACS2 (v2.2.9.1; broad mode, broad cut-off 0.1, q < 0.05). Merged peaks (BEDTools v2.31.1) present in both replicates were retained as high-confidence regions (16,525 primed; 9,377 unprimed). Genes whose gene body plus 2 kb upstream window overlapped a high-confidence region were considered targets and classified as P-only, UP-only, common or non-targets (36,140 genes). Signal was quantified as fragment Counts Per Million (CPM) per window with a library-size-scaled pseudocount, averaged across replicates and expressed as log2(primed/unprimed), used for ranking only.

Targets were integrated with expression data (**Table S1.1**; DEGs: FDR < 0.05, |log2FCh| ≥ 1.5). Primed log2FCh and DEG proportions of primed only (P-only) and unprimed only (UP-only) genes were compared using two-sided Mann–Whitney U and Fisher’s exact tests (Benjamini–Hochberg corrected; FDR). Paralog families were defined as connected components of an all-against-all DIAMOND blastp search (v2.2.6; e-value ≤ 1e−10, ≥ 40% identity, ≥ 50% coverage), and log2FCh of genes and their paralogs was compared using Spearman’s correlation. Enrichment of GO, GO slim, Pfam, InterPro, KEGG, Plant Reactome, MapMan, TF family and functional categories (TFs, stress response, HSPs, HSFs) was assessed by one-sided hypergeometric tests against annotated backgrounds (FDR < 0.05, ≥ 3 genes). Analyses were performed in Python (v3.9.21) and R (v4.3.3), and figures were generated with ggplot2 (v3.5.2).

### Statistical analysis

Phenotyping experiments were conducted with a minimum of three independent biological replicates unless stated otherwise. For gene expression comparisons in qRT-PCR assays, pairwise Student’s *t-*tests were performed using R. Differences in continuous and discrete variable distributions were assessed using the non-parametric Wilcoxon rank-sum test and Fisher’s exact test, respectively, with multiple testing corrections applied using the false discovery rate (FDR) method. To evaluate the enrichment of commonly differentially expressed genes (DEGs) between shoots and roots, a permutation test was conducted with 10,000 randomisations. In each iteration, the observed common DEGs were replaced with randomly selected expressed genes to generate a null distribution of overlap counts. Z-scores were computed based on the deviation of observed overlap from the mean of the randomised overlaps, and P-values were estimated as the fraction of permutations yielding an overlap equal to or greater than the observed value. All statistical analyses were performed using R. All tests were two-sided unless stated otherwise, and FDR < 0.05 were considered significant.

## Results

### Thermopriming enhances heat stress adaptability by streamlining transcriptional responses

To delineate how regulatory programmes are rewired during priming in crop plants, we examined the impact of thermopriming on the heat stress response of rice seedlings. We recently showed that exposure of naïve, unprimed five-day-old seedlings to 45 °C for 4 h led to growth impairment compared to the control (untreated seedlings) (Nair *et al*., 2026). To test the impact of thermopriming on the growth of rice seedlings, we pre-exposed the seedlings to a sub-lethal heat stimulus at 37 °C for 2 h (thermopriming), followed by a 2 h recovery period, and then subjected them to a growth impairing heat stress of 45 °C for 4 h (triggering heat stress) (**Figure 1a**). Measurements of shoot and root length and fresh weight (eight days post-triggering stress) depict better growth in primed compared to unprimed seedlings (**Figure 1b-d**). These findings establish that thermopriming aids better adaptability to acute heat stress during early seedling development in rice.

**Figure 1.**
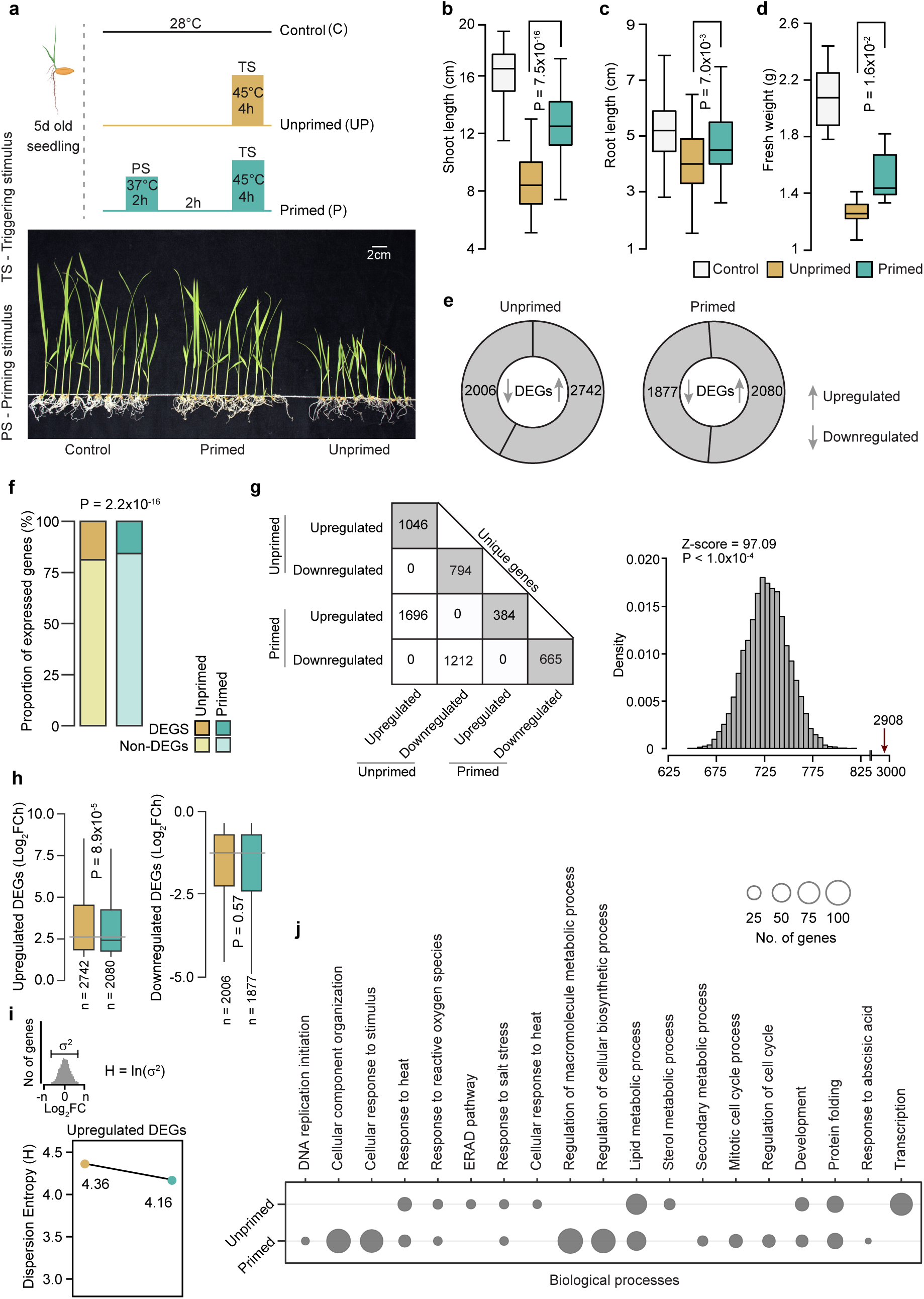
Thermopriming-induced transcriptional reprogramming confers heat stress tolerance. **(a)** Schematic representation of the experimental design. Five-day-old rice seedlings were either left untreated (control), exposed to a lethal triggering heat stress (unprimed), or subjected to a mild pre-treatment at 37 °C for 2 h and a recovery for 2 h (primed) before exposure to the triggering heat stress (45 °C for 4 h). Representative images showing shoot and root growth differences across control, primed, and unprimed seedlings eight days post-stress. Box plots indicating **(b)** shoot, **(c)** root length and **(d)** total seedlings’ fresh weight (*n =* 15 seedlings for all measurements; FW measurements were done by summation of the values for 15 seedlings together independently for three sets). *P* values were calculated using a two-tailed Student’s *t*-test. **(e)** Distribution of differentially expressed genes (DEGs) under unprimed and primed conditions following triggering heat stress. **(f)** Bar plot showing the proportion of DEGs and non-DEGs among the total expressed genes. *P* values were calculated using Fisher’s exact test. **(g)** Matrix plot showing the overlap of DEGs between unprimed and primed. Permutation test for checking the enrichment of DEGs overlap between unprimed and primed conditions. The grey histogram represents the random expectation, and the red arrow indicates the actual observation. **(h)** Boxplots showing the distribution of Log_2_ fold change values of the upregulated and downregulated DEGs. *P* values were calculated using the Wilcoxon rank sum test. **(i)** Line plot showing the dispersion entropy that was calculated based on the variance of gene expression distribution of upregulated DEGs. **(j)** Dot plot showing gene ontology (GO) enriched biological processes from DEGs in unprimed and primed seedlings. Dot size reflects the fold enrichment.

How does thermopriming confer such heat stress adaptability? To address this, we profiled the transcriptomes of control, unprimed, and primed rice seedlings after exposure to triggering heat stress (After triggering, AT). Surprisingly, we observed a significantly lower number of differentially expressed genes (DEGs) in response to stress in primed seedlings (3,957 DEGs; 2,080 upregulated and 1,877 downregulated) compared to unprimed seedlings (4,748 DEGs; 2,742 upregulated and 2,006 downregulated; **Figure 1e-1f; Table S1.1**). Nevertheless, both conditions shared a significant number of common DEGs (2,908 DEGs) (**Figure 1g**), alluding to an overlapping basic framework of heat stress response. In addition to a reduction in the number of DEGs, the magnitude of differential expression (represented by fold change), specifically for the upregulated DEGs, was significantly lower in primed than in the unprimed seedlings (**Figure 1h**). Estimation of dispersion entropy, which assesses the extent of variability in the differential expression within each condition (Zhu *et al*., 2020), revealed relatively lower entropy for upregulated DEGs in the primed condition compared to those of the unprimed condition (H=4.16 vs 4.36; **Figure 1i**). Notwithstanding a reduced number of DEGs, Gene Ontology biological processes (GO-BP) enrichment analysis revealed a higher number of enriched GO-BPs for DEGs upon priming compared to the unprimed state (**Figure 1j; Table S1.2**). Overall, thermopriming appears to elicit a streamlined transcriptional response that engages diverse stress-response processes, which may underlie improved adaptability of primed seedlings.

### Thermopriming-mediated transcriptional streamlining is potentially driven through constrained functional redundancy and H3K27me3-mediated gating

How is transcriptional streamlining achieved in primed seedlings? We posit that such streamlining could be driven through a systematic regulation of transcription factors (TFs) and/or epigenetic control. Indeed, primed plants showed fewer DETFs than unprimed plants (214 vs 285) across a similar number of TF families (**Figure 2a; Table S1.3**), with a lower magnitude of differential expression of upregulated DETFs (**Figure 2b**), and lower dispersion entropy among upregulated DETFs (H = 4.25 vs. 4.53; **Figure 2c**), compared to unprimed plants. Functional redundancies resulting from paralogs arising through gene-duplication (genetic redundancy), tend to immensely contribute during acute stress response (Zou *et al*., 2009). Interestingly, we find constrained genetic redundancies upon priming, wherein DETFs and DEGs in the primed condition are limited to small or moderately sized paralog clusters as against DETFs and DEGs in the unprimed condition, which tend to be preponderant in large paralog clusters (≥4 membership; **Figure 2d; Supplemental Figure 1; Tables S1.4 and S1.5**). Together, these point to a streamlined transcriptional response that could reduce functional redundancies or conflicts (Flavell, 1994; Garber *et al*., 2024) and the degree of gene-expression variability, thereby supporting a more efficient adaptive response upon priming.

**Figure 2.**
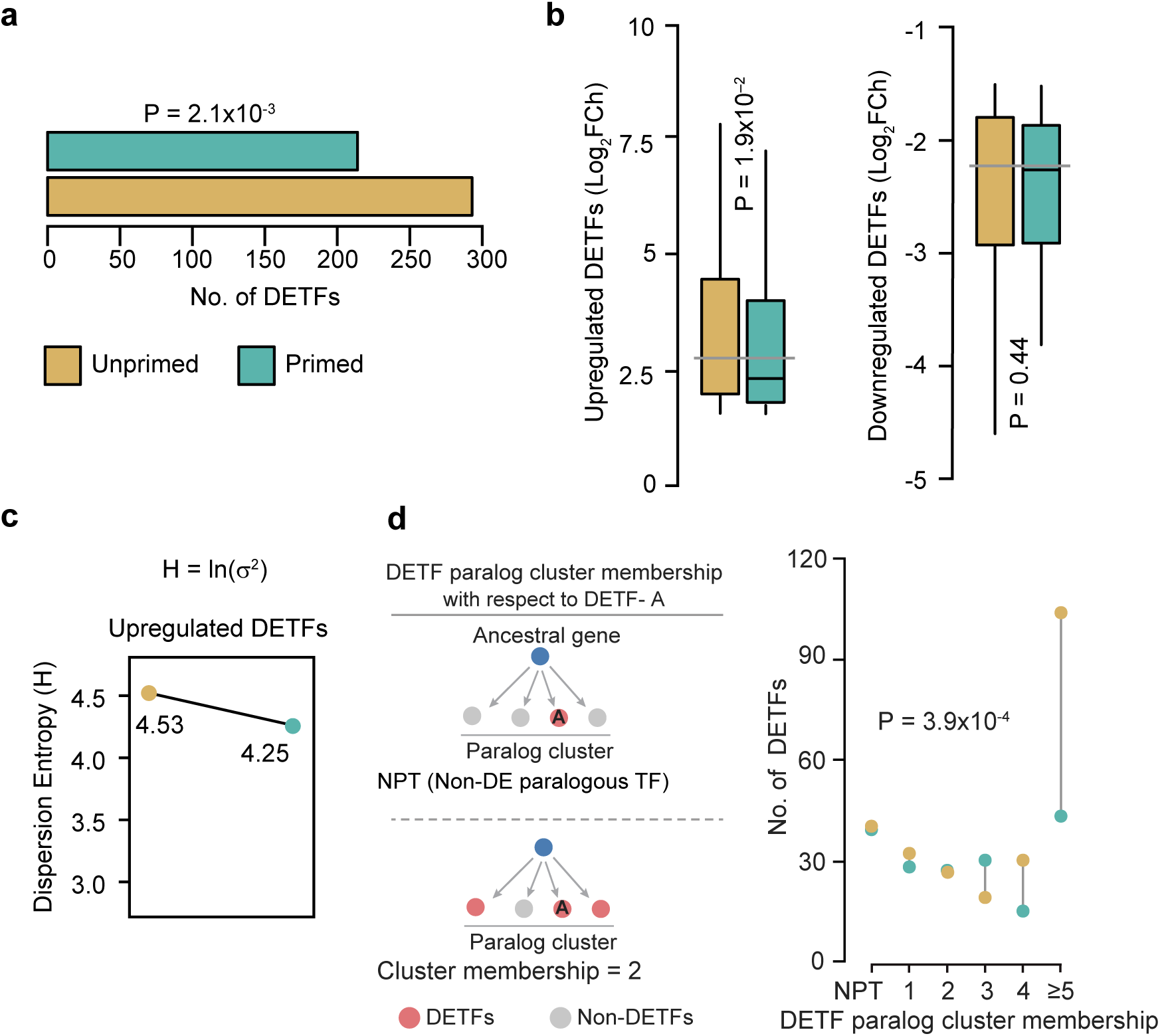
Thermopriming limits transcriptional regulatory redundancy by maintaining H3K27me3-mediated silencing. **(a)** The total number of DETFs in UP (n = 285) and P (n = 214). *P* values were calculated using Fisher’s exact test. **(b)** Boxplots showing the distribution of Log_2_ fold change values of the upregulated and downregulated DETFs. *P* values were calculated using the Wilcoxon rank sum test. **(c)** Dispersion entropy was calculated based on the variance of gene expression distribution for DE-TFs. **(d)** DEG paralog cluster size distribution for DETFs. *P* values were calculated using Fisher’s exact test.

What are the potential drivers of priming-mediated transcriptional streamlining? Given the shorter duration of priming-induced stress memory (2 h recovery) studied here, we hypothesised that the swift systematic alterations in gene expression could be primarily driven through reversible histone modifications that promote (active marks) or inhibit gene expression (repressive marks) (Lämke & Bäurle, 2017; Bäurle & Trindade, 2020; Ueda & Seki, 2020). In this regard, we probed for known histone marks (active: H3K27ac, H3K4me3 and repressive: H3K27me3) and a thermo-responsive histone variant H2A.Z whose eviction facilitates gene expression during heat stress (Kumar & Wigge, 2010), in control, primed control (exposed to only priming stimulus), primed and unprimed seedlings upon heat stress. We did not observe a difference in the total levels for the active marks H3K27ac, H3K4me3 and H2A.Z across all the conditions analysed. However, the repressive mark H3K27me3 was found to be significantly lower in unprimed seedlings compared to control or primed seedlings (**Figure 3a**).

**Figure 3.**
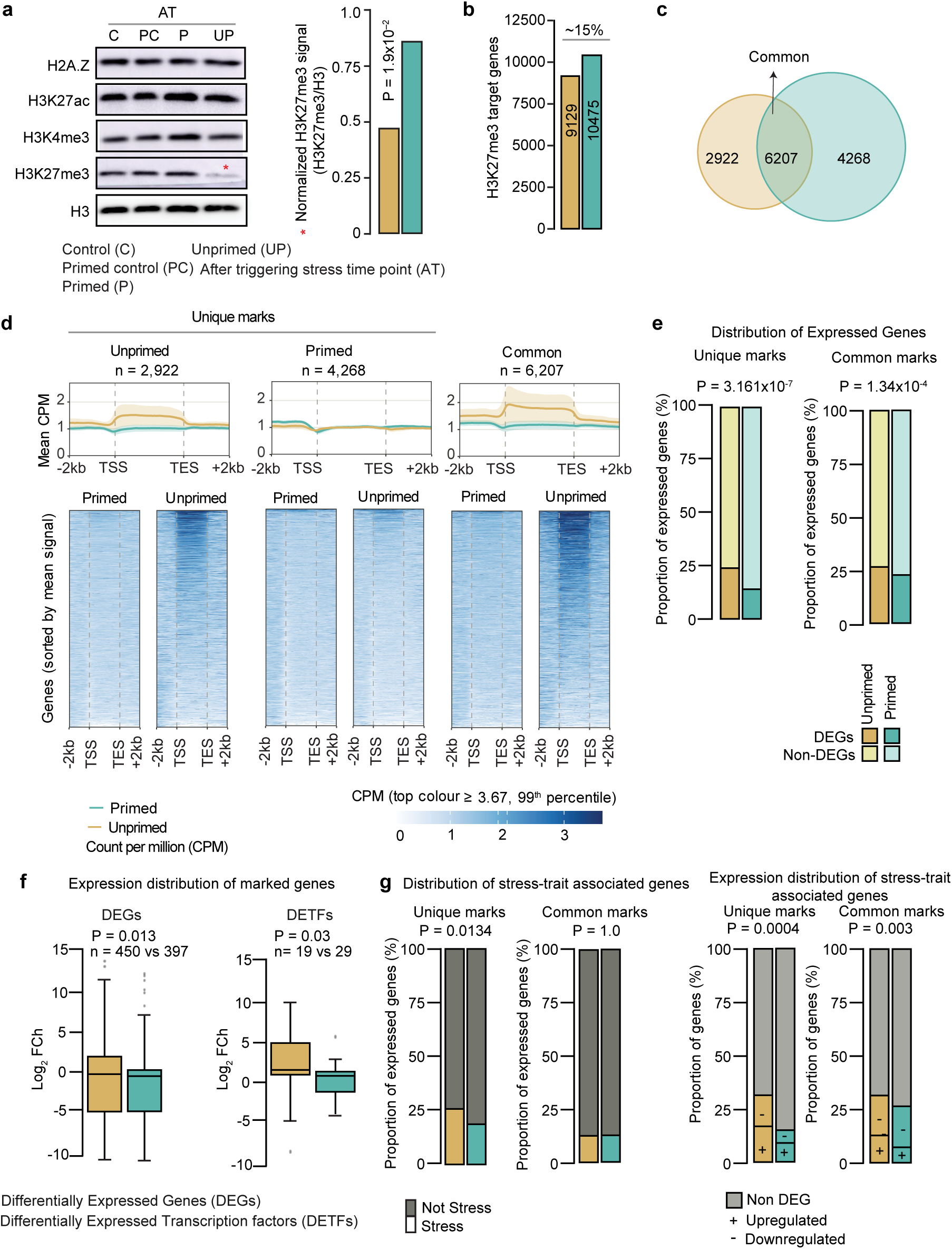
Retention of H3K27me3 upon priming contributes to the restrained transcriptional response of marked genes. **(a)** Western blot analysis of histone marks H3K27ac, H3K4me3, H3K27me3, and variant H2A.Z across control (C), primed control (PC), primed (P) and unprimed (UP) conditions after the triggering stress (AT); H3 serves as the loading control. Bar plot shows H3K27me3 signal normalised to H3 in unprimed and primed seedlings; the P value was calculated using a two-tailed Student’s t-test. ‘*’ indicates P < 0.05 (**b**) Number of H3K27me3-marked genes identified by ChIP-seq in primed and unprimed seedlings following triggering stress. **(c)** Overlap of H3K27me3-marked genes between unprimed (UP) and primed (P) seedlings. **(d)** Metagene profiles (top; mean CPM) and heatmaps (bottom) showing H3K27me3 signal across gene bodies, from 2 kb upstream of the transcription start site (TSS) to 2 kb downstream of the transcription end site (TES), for unprimed-only, primed-only and commonly marked genes. **(e)** Bar plots showing the proportion of expressed genes: differentially and non-differentially expressed genes (DEGs and non-DEGs) in uniquely and commonly marked genes in primed and unprimed. P value was calculated using the Fischer’s exact test. **(f)** Box plots showing the magnitude of expression changes for DEGs and DETFs marked in primed and unprimed seedlings. P values were calculated using the Wilcoxon rank sum test. **(g)** Bar plots showing the proportion of marked stress-related genes among uniquely and commonly marked genes (left; Fischer’s exact test), and those that were upregulated, downregulated or not differentially expressed (right; chi-square test).

To examine how H3K27me3 might gate transcription, we performed chromatin immunoprecipitation followed by sequencing (ChIP-seq) on primed and unprimed seedlings after triggering stress. Consistent with our immunoblots (**Figure 3a**), primed seedlings carried about 15% more H3K27me3-marked genes (10,475 vs. 9,129; **Figure 3b; Table S1.6**), including more condition-specific genes (4,268 vs. 2,922; **Figure 3c**). A large proportion of marked genes (6,207) was common between the two conditions, prompting us to ask whether the distribution of the marks along these genes is similar or different. Interestingly, unprimed seedlings showed elevated signal across gene bodies, whereas primed seedlings showed a broader set of marked genes with modest enrichment upstream of the transcription start site (TSS; **Figure 3d**). Thus, priming appears to spread H3K27me3 marking to more loci, preferentially at regulatory regions.

To relate H3K27me3 deposition to gene expression, we compared marked genes in primed and unprimed seedlings. This comparison revealed that H3K27me3 marking was associated with (i) significantly lower proportion of differentially expressed marked genes, from ∼24% (unprimed) to ∼14% (primed) for uniquely marked genes and from 26.5% to 22.6% for commonly marked genes (**Figure 3e**); and (ii) a dampened the magnitude of differential expression of uniquely marked DEGs and DETFs in the primed seedlings (**Figure 3f**). This restraint was stronger for genes that were previously reported to be stress-related (Oryzabase; https://shigen.nig.ac.jp/rice/oryzabase/gene/advanced/list). About 14.8% of marked stress-related genes were differentially expressed in primed seedlings compared with 31.4% in unprimed seedlings, and they made up a smaller share of marked DEGs (18.6% vs 25.8%; **Figure 3g**). Thus, H3K27me3 retention upon priming appears to limit both the expression of the number of genes in response to triggering stress and their magnitude, mirroring the streamlined transcriptional response (**Figures 1e–i and 2**).

We next asked whether H3K27me3 marking contributes to the constrained paralog redundancy observed upon priming (**Figure 2d; Supplemental Figure 1**). About 10% of differentially expressed paralogs unique to unprimed seedlings carried H3K27me3 uniquely in either primed or unprimed seedlings (**Supplemental Figure 2**). The position of the mark differed between conditions: in primed seedlings, marks were enriched in the upstream region (−2 kb of TSS; ∼60%), followed by the gene body (TSS to TES; ∼27%), whereas in unprimed seedlings, most marks spanned both regions (∼60%), with the remainder in the gene body (∼20%) or upstream region (∼20%) alone.

We then asked whether this differential marking could explain why these paralogs are differentially expressed only in unprimed seedlings. About 60% of paralogs carrying upstream marks in primed seedlings, where they were not differentially expressed upon priming, were upregulated in unprimed seedlings, where they lacked this mark. In contrast, paralogs marked at other positions tended to be downregulated in both conditions. Thus, upstream H3K27me3 retained upon priming may restrain the expression of these paralogs, whereas in unprimed seedlings, where the mark is less retained, other regulatory mechanisms may act alongside H3K27me3.

Collectively, these findings imply that constrained functional redundancy and transcriptional repression aided by repressive histone mark H3K27me3-mediated gating may jointly drive streamlining of gene expression in the primed seedlings.

**Supplemental Figure 1.**
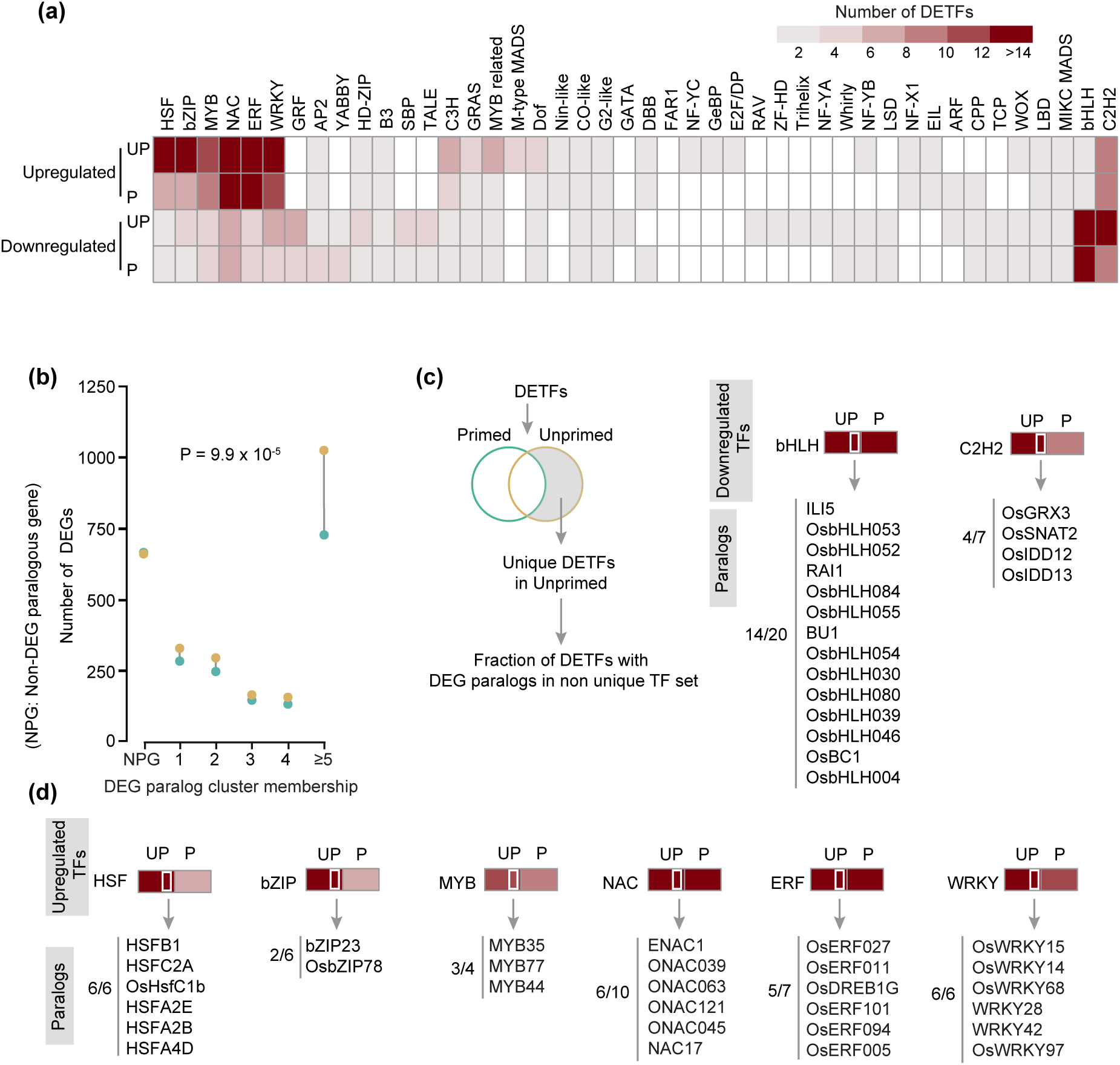
Paralog clusters of heat stress-specific DEGs and key transcription factors (TFs) in primed and unprimed conditions. (a) Heatmap showing the distribution of differentially expressed TF (DETFs) families in unprimed (UP) and primed (P) seedlings following triggering stress. **(b)** Distribution of differentially expressed gene (DEG) paralog cluster sizes under primed and unprimed conditions. *P* values were calculated using Fisher’s exact test. **(c)** Schematic outlining the identification of uniquely upregulated transcription factors (TFs) in unprimed seedlings that have DEG paralogs (left). Uniquely downregulated paralog TFs in unprimed seedlings for the bHLH and C2H2 TF families (right). **(d)** Uniquely upregulated paralog TFs in unprimed seedlings. Major TF families include HSF, bZIP, MYB, NAC, ERF, and WRKY.

**Supplemental Figure 2.**
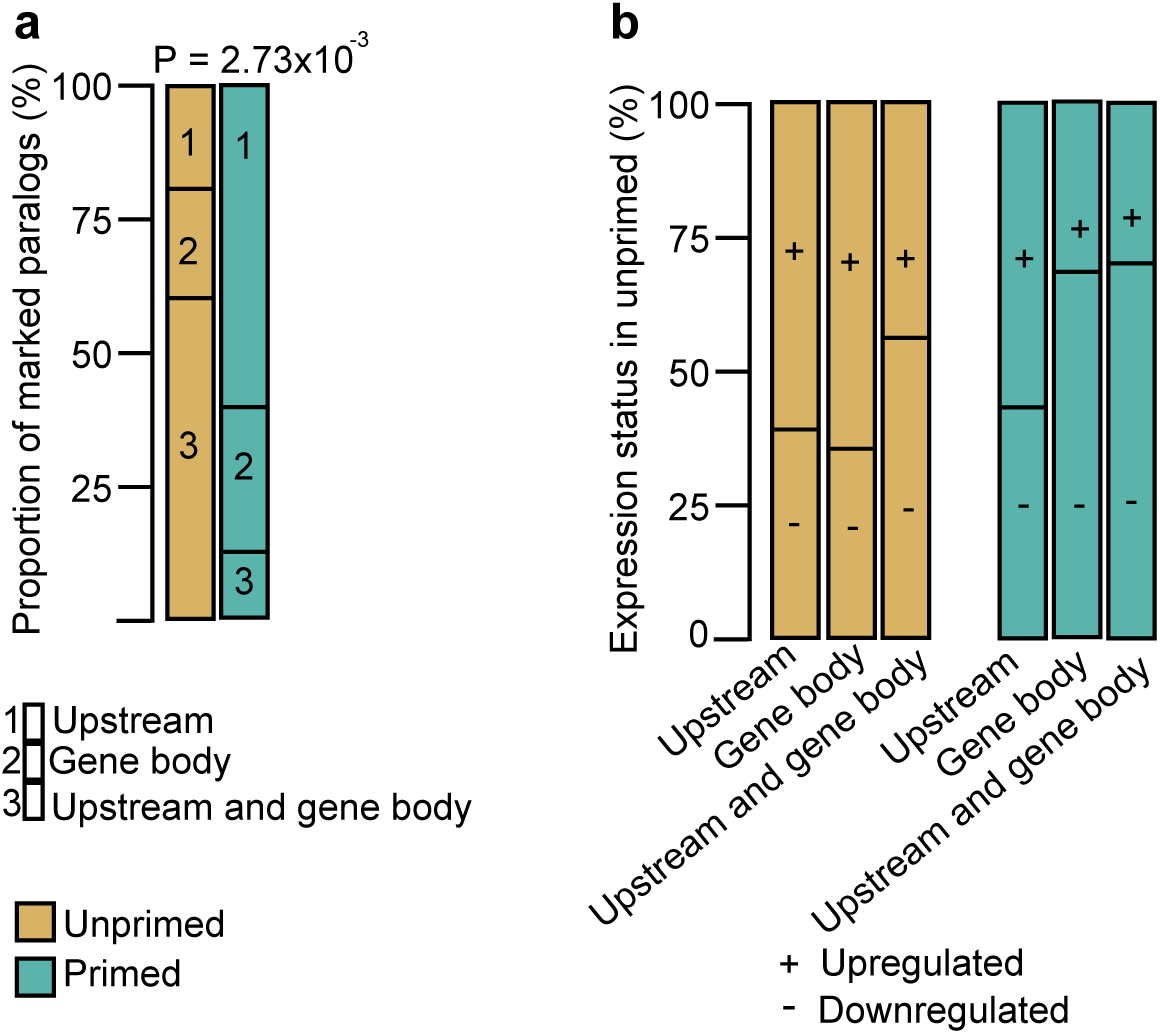
Effect of H3K27me3 marks on differently expressed paralogs unique to unprimed. (**a**) Bar plot shows the mark location of unprimed uniquely differentially expressed paralogs in primed (n= 81) and unprimed conditions (n= 83). P-value was calculated using the chi-square test. (**b**) Bar plot shows the expression distribution of marked genes correlative to different mark locations in primed and unprimed.

### Priming reduces regulatory network complexity while preserving growth-related modules

How does streamlined transcription aid in priming-mediated responses? To address this, we re-constructed the DETF-target regulatory networks as previously described by us (Nair *et al*., 2026) (**Figure 4a**). Briefly, putative target genes were identified by examining DEG promoter regions for enrichment of binding sites corresponding to DETFs. We then quantified the association between DETFs and their predicted targets using Pearson correlation analysis of their expression profiles. Subsequently, significant DETF-target pairs (with FDR<0.05) were used to reconstruct a high-confidence network of DEG targets potentially regulated by DETFs. Strikingly, the primed TF-target network was compact, consisting of 32 TFs and 983 targets with 1,808 interactions compared to 6,223 interactions among 46 TFs and 2,008 targets of the unprimed network. Not surprisingly, the number of targets regulated by each TF and the number of TFs regulating each target were significantly lower in primed seedlings compared to the unprimed seedlings (**Figure 4b**). Estimation of von Neumann entropy and Shannon entropy for the TF-target networks further confirmed a lower network entropy state in primed seedlings (**Figure 4c**). These observations underscore reduced transcriptional complexity in the primed condition.

**Figure 4.**
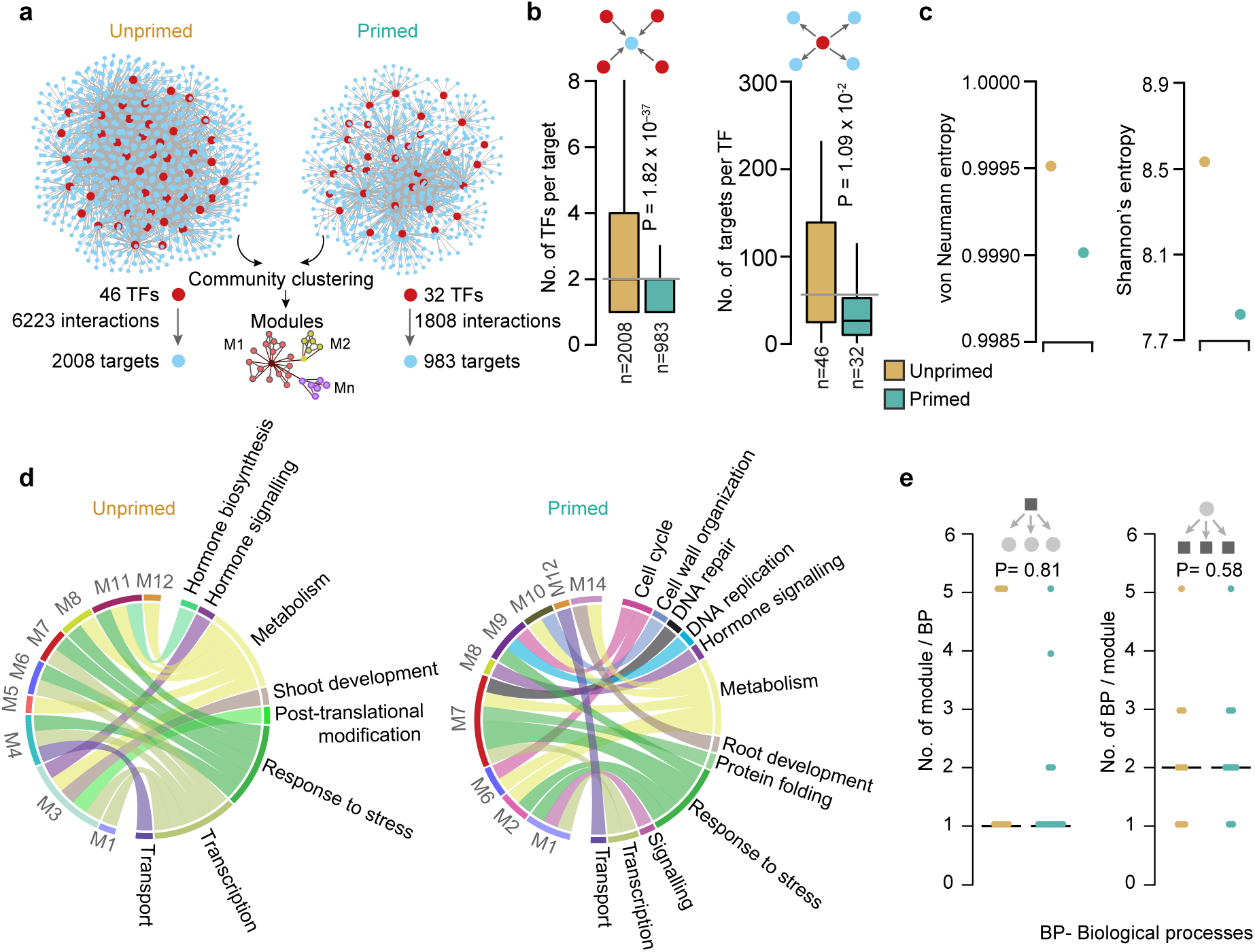
Thermopriming reprograms transcriptional regulatory networks to enhance specificity and reduce redundancy. **(a)** Network maps of inferred TF-target interactions in unprimed and primed seedlings. Nodes represent genes (red: TFs; blue: targets), and edges denote significant regulatory associations. **(b)** TF-target associations in the unprimed and primed network. *P* values were calculated using the Wilcoxon rank sum test. **(c)** Calculation of network entropy using von-Neumann and Shannon’s entropy. **(d)** Circos plots illustrating the biological processes regulated by unprimed and primed modules. **(e)** Quantitative analysis of the regulatory interactions between modules and biological processes in both unprimed and primed conditions. *P* values were calculated using the Wilcoxon rank sum test.

Next, we investigated the impact of reduced transcriptional complexity upon priming on the network architecture. For this, we analysed the primed and unprimed TF-target networks using the GLay community clustering algorithm (Su *et al*., 2010), which organises the TF-target networks into interconnected but distinct modules, as shown previously (Nair *et al*., 2026). Modularity in biological networks can contribute to efficient processing of information, increase robustness, adaptability, and reduce perturbations, collectively contributing to the fitness or survivability of organisms (Lorenz *et al*., 2011; Huitzil & Huepe, 2024). Interestingly, we found a comparable number of modulons between the primed (n=14) and the unprimed (n=12) conditions, though the primed TF-target network was considerably smaller in size. Akin to the network-level observations, modules of the primed TF-target network showed lower transcriptional complexity compared to that of the unprimed network (**Supplemental Figure 3a and 3b**). We next examined the enriched biological processes for each module across the two conditions and found that the distribution of biological processes across the modules and the number of modules regulating each biological process to be comparable (**Figure 4d and 4e; Table S1.7**). A closer look at the biological processes revealed common biological processes across primed and unprimed conditions, such as transport, transcription, response to stress, metabolism, and hormone signalling (**Figure 4d**). In addition, primed seedlings showed enrichment for distinct biological processes that regulate genetic information flow and contribute to growth, such as DNA replication (module M9), DNA repair (M8), protein folding (M7), cell wall organisation (M10), and cell cycle progression (M6 and M9). In contrast, the biological processes unique to the unprimed condition included shoot development and post-translational modification. These results suggest that priming preserves the regulation of biogenesis and maintenance processes, which may contribute to the better growth of primed seedlings (**Figure 1b–d**). Collectively, these findings indicate that orchestrated regulatory response during priming extends beyond reduced transcriptional complexity at the node-level. It also involves a well-defined modular organisation, where specific TF-regulated modules control distinct biological processes, which may enhance robustness and adaptability while sustaining growth.

**Supplemental Figure 3.**
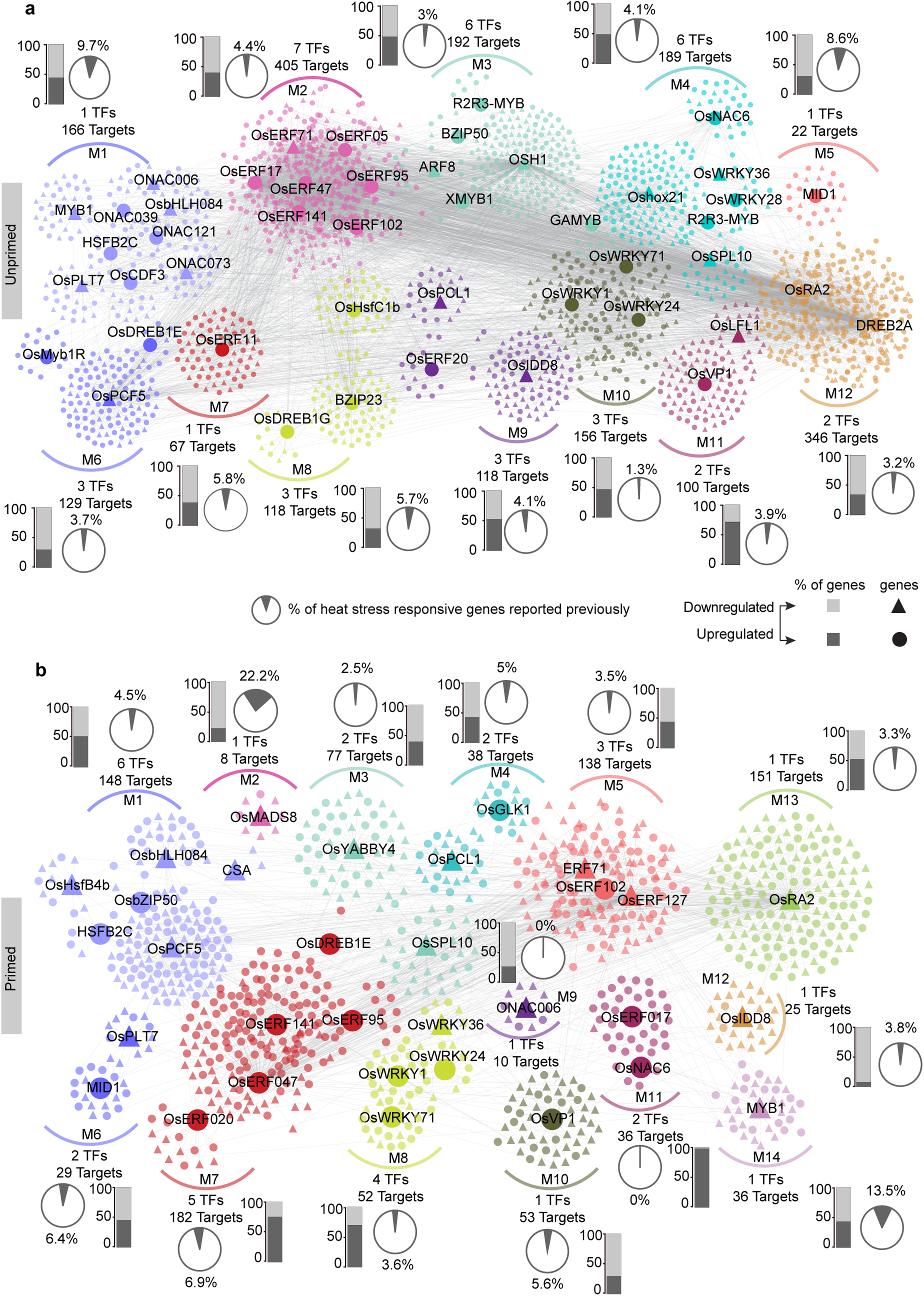
Modular organisation of transcription factor-target regulatory networks under heat stress. (a) TF-target network modules derived from unprimed seedlings. **(b)** TF-target network modules derived from primed seedlings. Modules represent groups of transcription factors (TFs) and their associated targets inferred from gene expression data. Bar plots adjacent to each module indicate the proportion of upregulated (black) and downregulated (grey) genes. Pie charts represent the fraction of genes within each module previously implicated in heat stress responses, based on the STIFDB database (Naika *et al*., 2013; https://caps.ncbs.res.in/stifdb2/).

### Priming-induced chaperone memory enables streamlined and effective heat shock response

Which priming-induced molecular changes contribute to the streamlined effective transcriptional response after triggering stress? We hypothesised that the transcriptional changes induced upon priming may contribute to the differential response of the primed seedlings upon triggering heat stress. To unveil this, we profiled the transcriptome changes immediately after exposure to priming (after priming, AP) relative to respective control seedlings (**Figure 5a**). At AP, 200 genes were differentially expressed (112 upregulated and 88 downregulated genes), which showed enrichment for the biological processes ‘protein folding’ and ‘response to heat’ (**Figure 5b and 5c**). Analysis of protein classes using protein domain-based annotations for DEGs at AP revealed chaperones to be the most predominant class (**Figure 5d**). These results suggest that the enhanced levels of chaperones after priming could drive better heat stress response in primed seedlings after triggering stress.

**Figure 5.**
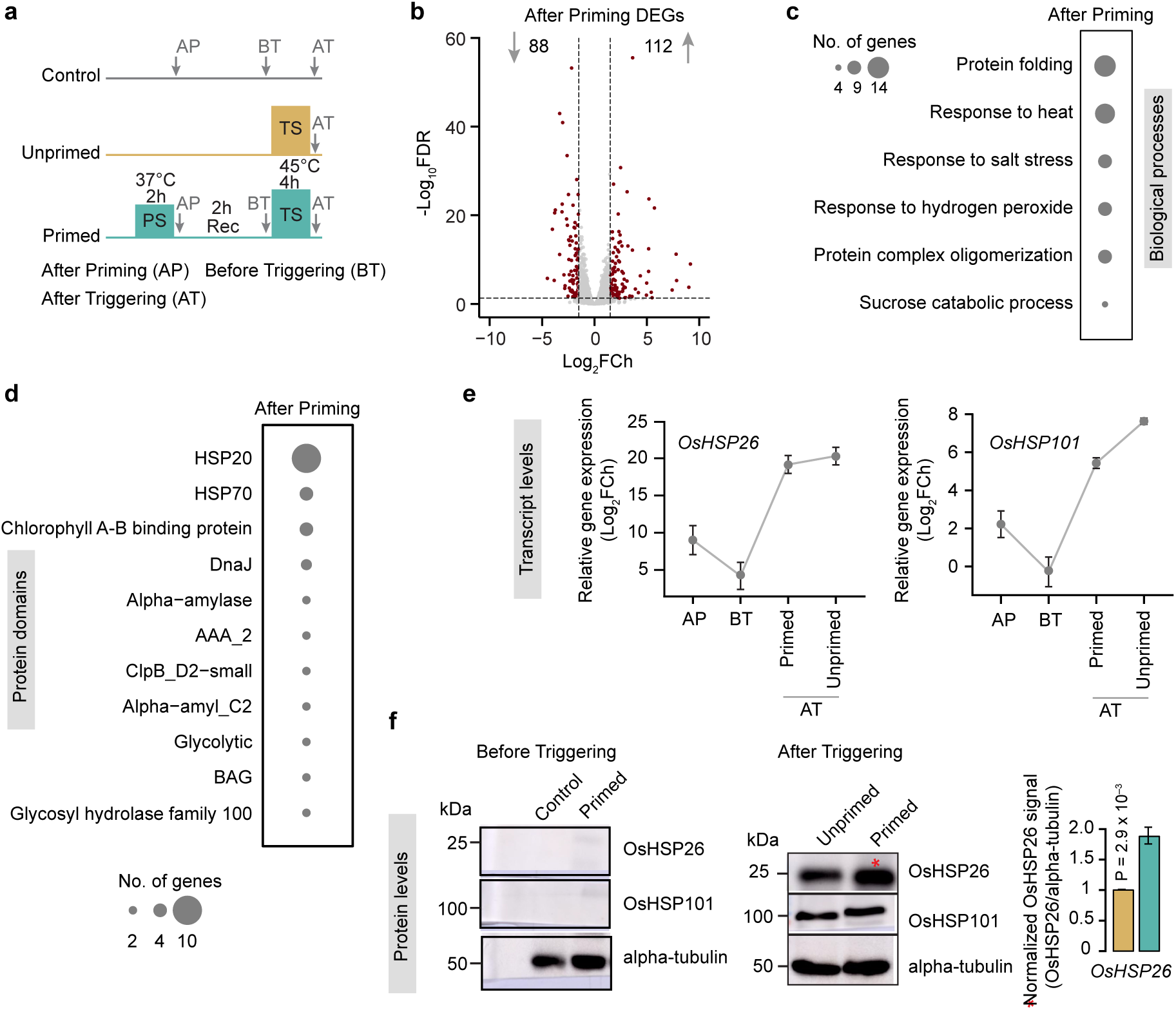
Priming-induced chaperone memory enables a streamlined yet effective heat stress response. **(a)** Schematic of the experimental design. Sampling time points include after priming (AP), before triggering (BT), and after triggering (AT). **(b)** Volcano plot showing differentially expressed genes (DEGs) in primed plants at the after-priming (AP) time point relative to the control. DEGs were identified with thresholds of Log2fold change ≥1.5 or ≤ −1.5 and false discovery rate (FDR) <0.05. DEGs are shown in brown. Numbers indicate the total count of significantly downregulated (88) and upregulated (112) genes. **(c)** Bubble plot of enriched biological process Gene Ontology (GO) terms among DEGs in primed plants after priming. Bubble size corresponds to the number of genes assigned to each term, as indicated in the legend. **(d)** Bubble plot showing enriched protein domain categories among DEGs in primed plants after priming. Bubble size corresponds to the number of genes, as indicated. **(e)** Relative transcript levels (Log₂FCh) of *OsHSP26 (left) and OsHSP101* (right) across time points (AP, BT and AT) and conditions (primed and unprimed). Data represent mean±S.E. **(f)** Immunoblot analysis of OsHSP26, OsHSP101, and alpha-tubulin (loading control) protein levels before triggering (left; control versus primed) and after triggering (right; unprimed versus primed). Molecular weights (kDa) are indicated. The bar graph (right) shows quantification of OsHSP26 signal normalised to alpha-tubulin in primed and unprimed plants after triggering. Data represent mean±S.E. *P* values (indicated by ‘*’) were calculated using two-tailed Student’s *t*-tests.

To test this, we investigated the expression profiles of the rice orthologs of a well-characterised Arabidopsis heat stress memory chaperone, AtHSP21 (rice ortholog: OsHSP26; Sedaghatmehr *et al*., 2016) and the non-memory heat stress-responsive chaperone, AtHSP101 (OsHSP101; Brzezinka et al., 2016; Lämke et al., 2016). As the primed response to triggering stress relies on heat stress memory, we compared the temporal expression dynamics of the heat stress memory chaperone with the non-memory but heat stress-responsive chaperone. At AP time point, both genes showed induced expression (**Figure 5e**). At the before triggering (BT) time point (end of recovery phase), *OsHSP26* transcripts remained slightly elevated, whereas *OsHSP101* returned to levels seen in the control seedlings (**Figure 5e**). After triggering stress (AT), transcript levels of both chaperones were slightly lower in primed than in unprimed seedlings (**Figure 5e**). We next asked whether the memory chaperone is also retained at the protein level at BT. However, protein levels of both HSPs were comparable between primed and control seedlings at BT (**Figure 5f**). In contrast, after triggering stress, primed seedlings accumulated significantly more OsHSP26 protein than unprimed seedlings, whereas OsHSP101 levels appeared similar (**Figure 5f**). Thus, despite slightly lower transcript levels at AT, primed seedlings accumulate more memory chaperone protein, suggesting that priming-mediated retention of OsHSP26 transcripts during recovery facilitates its enhanced accumulation upon triggering stress. Notably, primed seedlings achieved higher memory chaperone protein levels with lower transcriptional investment at AT, mirroring the reduced number and magnitude of DEGs observed upon triggering stress (**Figure 1e–j**). Thus, priming appears to pre-wire the protein maintenance machinery, allowing primed seedlings to mount an effective heat shock response without the transcriptional burden seen in unprimed seedlings, which may underlie their better growth after triggering stress (**Figure 1b–d**).

## Discussion

Thermomemory models, thus far, are largely based on boosting/re-induction of individual memory genes and their regulators, such as HSFs and chromatin modifiers, upon repeated stress (Lämke *et al*., 2016; Friedrich *et al*., 2021; Yamaguchi *et al*., 2021; Nair *et al*., 2022). Here we take a new approach to priming-mediated stress responses in a crop plant, investigating how transcriptional reprogramming after priming shapes the response to repeated stress, by integrating global analysis of different layers of genetic information flow in a time-resolved manner. The findings presented here highlight that primed seedlings mount a streamlined transcriptional response to triggering stress, indicating that memory both restrains and amplifies the response. This restraint is associated with a distinct regulatory architecture: fewer responding transcription factors (TFs) from smaller paralog families, more compact TF–target networks and retention of the repressive histone mark (**Figure 6**), suggesting a regulatory state pre-wired upon priming.

**Figure 6.**
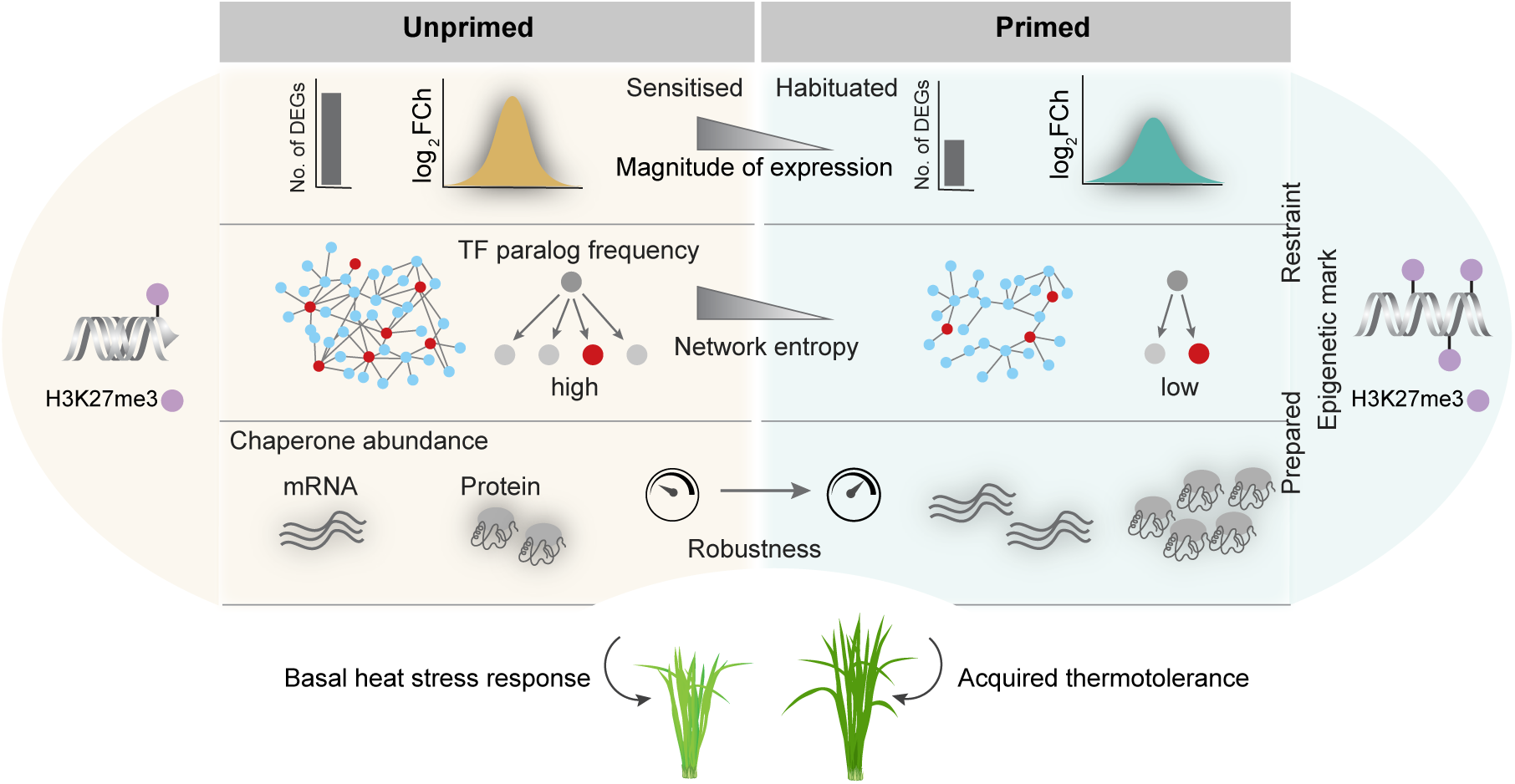
Proposed model of how thermopriming restrains and prepares the transcriptional response to triggering heat stress. A model showing the difference in molecular regulation in the unprimed and primed seedlings in response to acute heat stress. Unprimed seedlings mount a broader transcriptional response, with more DEGs, a larger TF–target network and greater disorderliness, which is associated with reduced growth after triggering stress (Figure 1b**–d**). By contrast, primed seedlings mount a smaller, more ordered response, involving fewer TFs drawn from smaller paralog families and a more compact network, and associated with retention of H3K27me3 at more genes, which are less likely to respond to stress. Together with greater accumulation of the memory chaperone OsHSP26 (Figure 5e**–f**), these features are proposed to support heat stress tolerance with a smaller growth penalty.

Acquired thermotolerance involves sustained or enhanced re-induction of stress-responsive genes (Lämke *et al*., 2016; Friedrich *et al*., 2021). However, prolonged stress responses may impair growth and are thought to be actively reset (Crisp *et al*., 2016; Sedaghatmehr & Balazadeh, 2024; Staacke *et al*., 2025). Interestingly, we find that thermopriming shapes a lean transcriptional response, enabling a swift, energetically efficient and robust heat stress response. Priming results in fewer differentially expressed genes, with modest expression changes, lower entropy and a compact TF–target network, together with improved growth after triggering stress. Similar patterns have been observed in other systems. In bacteria, higher transcriptional entropy predicts poor fitness under stress (Zhu *et al*., 2020), and drought-tolerant *Brassica napus* genotypes show a more restrained, targeted response than sensitive ones (Huang *et al*., 2026).

Priming may be favoured where stress recurs predictably and is more economical than repeatedly mounting a full defence response (van Hulten *et al*., 2006; Mitchell *et al*., 2009; Hilker *et al*., 2016). We find that priming induces the expression of TFs from smaller paralog families, whereas the induction of expression of large, duplicated families, as seen in unprimed seedlings, typically drives the acute stress response (Hanada *et al*., 2008; Zou *et al*., 2009; Jiang *et al*., 2013). This suggests that priming routes a restrained response through a core set of less redundant regulators. Further, our findings point to a complementary role for retained H3K27me3, a repressive histone mark, in restraining gene expression. Priming-mediated retention of H3K27me3 may therefore limit arbitrary gene activation, whereas its removal in unprimed seedlings coincides with an expansive response. This is consistent with, rather than contrary to, H3K27me3 removal at memory genes (Yamaguchi & Ito, 2021; Yamaguchi *et al*., 2021), suggesting a complementary strategy in which demethylation activates a select gene set and retention elsewhere tempers the wider heat stress response. Although causality remains untested, the association of H3K27me3 retention and restrained transcriptional response offers a new view of epigenetic regulation of plant resilience. Such restraints may therefore be beneficial where heat episodes recur within hours to days.

What might drive the restrained transcriptional response upon priming, in contrast to a broader response in unprimed seedlings? Disrupted proteostasis, a major problem for cells under heat stress, could drive more stress-responsive genes to be expressed at higher magnitude (Sugio *et al*., 2009; Masser *et al*., 2019). Priming may counter this through transcriptional activation of protein quality control mechanisms, along with retention of memory chaperone transcripts, such as OsHSP26, which together lead to higher accumulation of chaperone proteins upon the triggering stress. This could preserve proteostasis and keep proteins functional, protecting them from heat-induced unfolding, misfolding and degradation. Such robust proteostasis might in turn facilitate a more streamlined regulatory reprogramming in primed seedlings, with fewer differentially expressed genes, less functional redundancy, and lower expression magnitude.

Together, these observations point to a phenomenon in which thermopriming pre-wires the response to recurring heat by establishing a regulatory state that both prepares and tailors the later response, both during priming and recovery (**Figure 6**). Preparation is reflected in the chaperone-mediated preservation of proteostasis and restraint is reflected by the expression of a limited set of stress-responsive regulators, possibly aided by retention of the repressive histone mark H3K27me3.

Whether this pre-wired transcriptional reprogramming is conserved across species remains to be determined. Other mechanisms, not examined here, may also contribute to priming-mediated preparation and restraint, including negative feedback on HSF activity by accumulating chaperones, promoter architecture, control of RNA polymerase II activity at memory genes (Crawford *et al*., 2024) and post-transcriptional regulation such as mRNA turnover and alternative splicing (Crisp *et al*., 2016). Thus, the coordinated, pre-wired regulatory networks that result from priming may reflect efficient adaptation to recurrent heat in specific rice-growing environments and could be mechanisms that have been selected for in heat-resilient crop varieties.

## Acknowledgements

ADA thanks the USDA-FAS Norman E Borlaug Award for the financial support to conduct the RNA sequencing experiments. ADA and FBF thank partial funding provided by USDA-FAS project # FX22SR-10961R006. SV thanks the University Grants Commission (UGC), Government of India, for the doctoral fellowship. We thank IISER Tirupati core funding (ADA, SC, RVK, AUN), Ramalingaswami Re-entry Fellowship (BT/RLF/Re-entry/05/2018) from Department of Biotechnology, Government of India (SC), Core Research Grant (CRG/2023/004691) from Anusandhan National Research Foundation, Government of India (SC).

## Author contributions

ADA conceived the idea and supervised the work. AUN performed the phenotyping and immunoblot, ChIP-seq experiments and RNA-seq data analysis. RVK and SV performed the ChIP-seq analysis and relevant figure generation. AKS prepared the initial draft. ADA and SC wrote and edited the manuscript. ADA, SC, AUN, SV were involved in designing the computational analysis, generation of figures, interpreting the data. FBF was involved in editing the manuscript.

## Data availability

The raw transcriptome data are submitted to the Sequence Read Archive (SRA) of the National Centre for Biotechnology Information (NCBI) and will be made available upon the acceptance of the manuscript.

